# All-trans retinoic acid induces GADD34 gene expression via transcriptional regulation by Six1-TLE3 and post-transcriptional regulation by p38-TTP in skeletal muscle

**DOI:** 10.1101/2021.10.04.463012

**Authors:** Yuichiro Adachi, Masashi Masuda, Iori Sakakibara, Takayuki Uchida, Yuki Niida, Yuki Mori, Yuki Kamei, Yosuke Okumura, Hirokazu Ohminami, Kohta Ohnishi, Hisami Yamanaka-Okumura, Takeshi Nikawa, Yutaka Taketani

**Affiliations:** Department of Clinical Nutrition and Food Management, Institute of Biomedical Sciences, Tokushima University Graduate School, Tokushima, Tokushima 770-8503, Japan; Department of Nutritional Physiology, Institute of Biomedical Sciences, Tokushima University Graduate School, Tokushima, Tokushima 770-8503, Japan

**Keywords:** MAPK, mRNA stabilization, skeletal muscle type change, transcription, unfolded protein response

## Abstract

All-trans retinoic acid (ATRA) increases the sensitivity to unfolded protein response (UPR) in differentiating leukemic blasts. The downstream transcriptional factors of PERK, a major arm of UPR, regulates muscle differentiation. However, the role of growth arrest and DNA damage-inducible protein 34 (GADD34), one of the downstream factors of PERK, and the effects of ATRA on GADD34 expression in muscle remain unclear. In this study, we identified ATRA increased the GADD34 expression independent of the PERK signal in the gastrocnemius muscle of mice. ATRA up-regulated GADD34 expression through the transcriptional activation of it via inhibiting the interaction of homeobox Six1 and transcription co-repressor TLE3 with the MEF3-binding site on the *GADD34* gene promoter in myoblasts. ATRA also inhibited the interaction of TTP, which induces mRNA degradation, with AU-rich element on *GADD34* mRNA via p38 MAPK, resulting in the instability of *GADD34* mRNA. Overexpressed GADD34 in myoblasts changes the type of myosin heavy chain in myotubes. These results suggest ATRA increases GADD34 expression via transcriptional and post-transcriptional regulation in myoblasts, which changes muscle fiber type in myotubes.

## Introduction

The endoplasmic reticulum (ER) is a membranous organelle that has a central role in protein biosynthesis. Various stresses cause the accumulation of unfolded proteins in the ER, which induces ER stress (Walter & Ron, 2011; Szegezdi *et al*, 2006). An excessive ER stress response can result in various diseases, such as diabetes, inflammation, and cardiovascular diseases, including vascular calcification (Kim *et al*, 2008; Masuda *et al*, 2013; Masuda *et al*, 2015). To alleviate ER stress, the unfolded protein response (UPR) is initiated by the activation of 3 ER transmembrane sensors: PKR-like endoplasmic reticulum kinase (PERK), inositol-requiring enzyme 1 (IRE1), and activating transcription factor 6 (ATF6). PERK activation leads to the phosphorylation of a subunit of eukaryotic initiation factor 2 (eIF2α), resulting in the upregulation of activating transcription factor 4 (ATF4) and C/EBP-homologous protein (CHOP).

UPR in skeletal muscle regulates muscle stem cell homeostasis and myogenic differentiation. Especially, the PERK pathway is one of the key signals to promote the progression and commitment of satellite cells to the myogenic lineage (Rayavarapu *et al*, 2012; Gallot *et al*, 2019). Deletion of *PERK*, *ATF4*, or *CHOP* in satellite cells inhibits myofiber formation via the decreased expression of myoblast determination protein 1 (MyoD) and myogenin, essential transcription factors to correctly differentiate in skeletal muscle following injury (Gallot *et al*, 2019; Alter & Bengal, 2011; Ebert *et al*, 2020). We recently reported that UPR is an important regulator of injured skeletal muscle of mice with chronic kidney disease (Niida *et al*, 2020). Furthermore, sine oculis homeobox homolog 1 (Six1), is a mandatory transcription factor with coactivator eyes absent (Eya) and dachshund (Dach) or corepressor Groucho/transducing-like enhancer of split (TLE) for myogenic differentiation (Jennings & Horowicz, 2008; Sakakibara *et al*, 2016; Maire *et al*, 2020). However, little is known about the relationship between Six1 and PERK-related gene in muscle differentiation.

ATF4 promotes the transcriptional activation of growth arrest and DNA damage-inducible protein 34 (GADD34/Ppp1r15a), which dephosphorylates eIF2α by interacting with the catalytic subunit of type 1 protein serine/threonine phosphatase (PP1) (Harding *et al*, 2000; Zebrucka *et al*, 2016). Therefore, ATF4-mediated induction of GADD34 functions as a negative feedback loop of UPR by eIF2α dephosphorylation, which is essential for cell survival. Although GADD34 plays various roles without UPR signaling in some cells, such as cytokine production in dendritic cells, inhibition of apoptosis in liver cancer cells, and the response to chronic oxidative stress in neurodegenerative diseases (Clavarino *et al*, 2012; Song *et al*, 2019; Goh *et al*, 2018), the specific role of GADD34 in skeletal muscle has not been revealed.

All-trans retinoic acid (ATRA), a metabolite of vitamin A, is a ligand for nuclear receptors, including retinoic acid receptors (RARα, β, and γ). RARs in the nucleus work as a transcription factor by forming a heterodimer with retinoid X receptors (RXRs) and binding to retinoic acid-response elements (RAREs) in target gene promoters (Chambon, 1996; Mangelsdorf & Evans, 1995). ATRA also activates various MAPK kinase cascades, such as p38 and ERK (p44/42), with RARα in membrane lipid rafts and RARγ in the cytosol (Eochette-Egly, 2015). In addition, activated p38 and ERK are involved in not only the transcriptional regulation of the target gene via translocation of the target gene to the nucleus but also the post-transcriptional regulation of the target mRNA by interacting with the mRNA binding proteins (RBPs), such as Tristetraprolin (TTP), which controls mRNA stability (Bhattacharyya *et al*, 2011; Stoecklin *et al*, 2004). ATRA has an important role in the differentiation and apoptosis of blood cells and the sensitivity to ER stress through the differentiation of human leukemic cells (Masciarelli *et al*, 2018). In contrast, ATRA supplementation ameliorates the ethanol-induced expression of ATF4 and CHOP in the rat liver (Nair *et al*, 2018). Interestingly, ATRA also controls muscle differentiation via the transcriptional regulation of MyoD and myogenin (Chen *et al*, 2015; Lamarche *et al*, 2015), whereas the effects of ATRA on PERK-related genes in skeletal muscle cells is still unknown.

In this study, we investigated the effects of ATRA on the PERK pathway in various organs, especially muscle. Our results demonstrate that ATRA increases the *GADD34* gene expression via the downregulation of Six1 and the stabilization of *GADD34* mRNA in skeletal muscle. We also characterized the role of GADD34 in skeletal muscle fiber type change.

## Results

### ATRA induces *GADD34* gene expression in the gastrocnemius muscle and undifferentiated skeletal muscle cells without translational activation of ATF4

To assess the response of the PERK signal to ATRA in various tissues, such as the kidney, liver, epididymal adipose (EA), gastrocnemius muscle (GM), and femur, mice were treated with ATRA (total dose: 10 mg/kg body weight). The expression of *ATF4* and *CHOP* mRNA were not changed in all tissues (as described above) of mice treated with ATRA for 24 h. ATRA treatment significantly increased the *GADD34* mRNA expression only in the GM of mice (Fig 1A). Western blotting revealed that ATRA treatment did not increase phosphorylated eIF2α (p-eIF2α), ATF4, or CHOP downstream of the PERK signal in GM in comparison to vehicle; however, it increased GADD34 protein expression levels (Fig 1B). To determine whether the increase in the expression of GADD34 by ATRA is associated with the differentiation of skeletal muscle cells, C2C12 cells were gradually differentiated from myoblasts to myotubes, as shown in Fig 1C. In C2C12 myoblasts, myotubes, and HEK293 cells, the expression of *ATF4* mRNA was comparable between DMSO and ATRA treatment, which was in line with the results of *in vivo* studies. Surprisingly, ATRA increased the GADD34 mRNA and protein expression levels in C2C12 undifferentiated myoblasts, but not in myotubes or HEK293 cells (Fig 1D and E). We generated C2C12 cells with the knockdown of ATF4 to confirm the effects of ATRA on the GADD34 expression independently of ATF4; with the use of ATF4-specific siRNA, the endogenous *ATF4* mRNA levels of these C2C12 cells were reduced by more than 60%. ATF4-knockdown did not inhibit the induction of the expression of *GADD34* mRNA by ATRA (Fig 1F).

**Figure 1.**
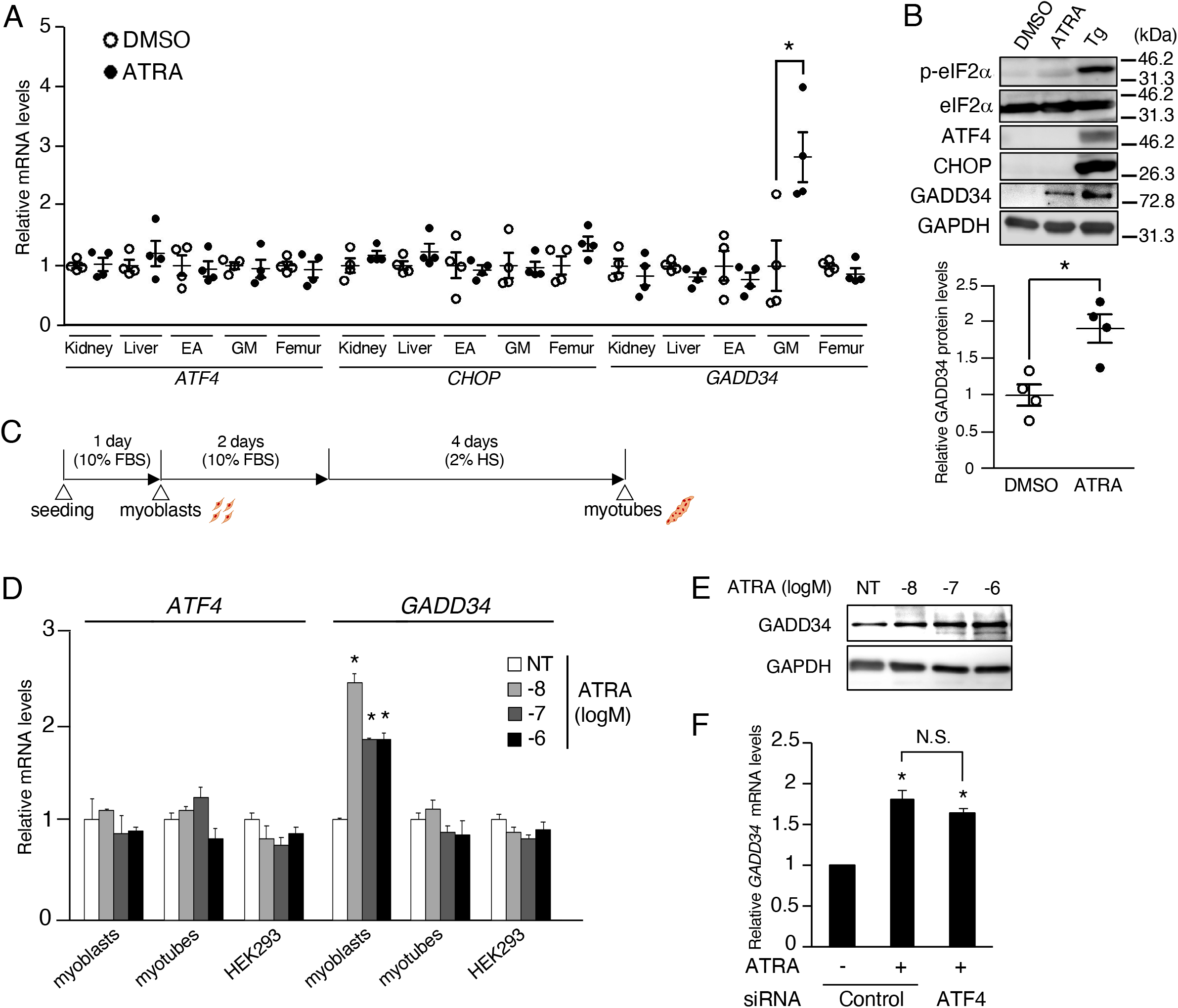
Effects of ATRA treatment on PERK signaling in skeletal muscle. A, B Eight-week-old male C57BL/6J mice were given an intraperitoneal injection of 0.1% DMSO (vehicle) and ATRA (10 mg/kg) for 24h. (A) The mRNA expression levels of *ATF4*, *CHOP*, and *GADD34* in kidney, liver, epididymal adipose (EA), gastrocnemius muscle (GM), and femur of mice were determined by real-time PCR (*n* = 4 mice per group). (B) Western blotting of phosphorylated-eIF2α (p-eIF2α), total eIF2α (eIF2α), ATF4, CHOP, and GADD34 in GM (*n* = 4 mice per group). C Schematic illustration of the experimental timeline in C2C12. D The mRNA expression levels of *ATF4* and *GADD34* in C2C12 cells (myoblasts and myotubes) and HEK293 cells treated with the indicated concentrations of ATRA (*n* = 3–4). E Western blotting of GADD34 in C2C12 cells myoblasts with the indicated concentrations of ATRA. F The mRNA expression levels of *GADD34* in C2C12 myoblasts transfected with *ATF4* siRNA (siATF4) or control and incubated in the presence of 1 μM ATRA or DMSO as a vehicle for 24 h (*n* = 3–4). Data information: In (A–B), data are presented as the mean ± SEM. \**p* < 0.05 (two-tailed unpaired Student’s *t*-test). In (D, F), data are presented as the mean ± SEM. \**p* < 0.05 vs. DMSO (NT) (one-way ANOVA with a Student-Newman post-hoc test). N.S. = not significant. Source data for this figure are available online.

### ATRA upregulates the transcriptional activity of the *GADD34* gene in C2C12 cells

To investigate the molecular mechanisms underlying the undifferentiated muscle-specific regulation of the *GADD34* gene expression by ATRA, we examined the responsiveness of human *GADD34* gene promoters to ATRA using a luciferase assay with undifferentiated C2C12 myoblasts and HEK293 cells. The overexpression of RARs increased the luciferase activity of pGADD34-0.5k in both C2C12 myoblasts and HEK293 cells. ATRA increased the luciferase activity of pGADD34-0.5k in C2C12 myoblasts expressing RARα and RARγ, but not RARβ (Fig 2A). ATRA dose-dependently increased the *GADD34* gene promoter activity in C2C12 myoblasts, unlike in HEK293 cells (Fig 2B). Surprisingly, TTNPB, a major agonist of RAR, did not increase the *GADD34* gene promoter activity, although TTNPB increased the mRNA expression of the target gene (Appendix Fig S1A and B) (Thé *et al*, 1990). Unlike TTNPB, ATRA can activate the non-genomic p38 and ERK MAPK signals via extranuclear RARα/γ and RARγ, respectively (Tanoury *et al*, 2013; Bouchard & Paquin, 2013; Khatib *et al*, 2019). Next, we tested whether these MAPK signals are involved in the regulation of the GADD34 expression by ATRA. As a result, the ATRA-induced *GADD34* gene promoter activity was blocked by p38 inhibitor SB203580 or ERK inhibitor FR180204 via RARα/γ or RARγ, respectively (Appendix Fig S1C). Next, to explore the muscle-specific regulatory element of ATRA on the human *GADD34* gene promoter, several reporter constructs lacking portions of the 5’-promoter region of the human *GADD34* genes were tested in C2C12 myoblasts overexpressing RAR/RXR, with or without ATRA. These deletion analyses suggest that the sequence from -162 to -131 is responsible for ATRA-dependent activation of the human *GADD34* gene promoter activity in C2C12 myoblasts (Fig 2C).

**Figure 2.**
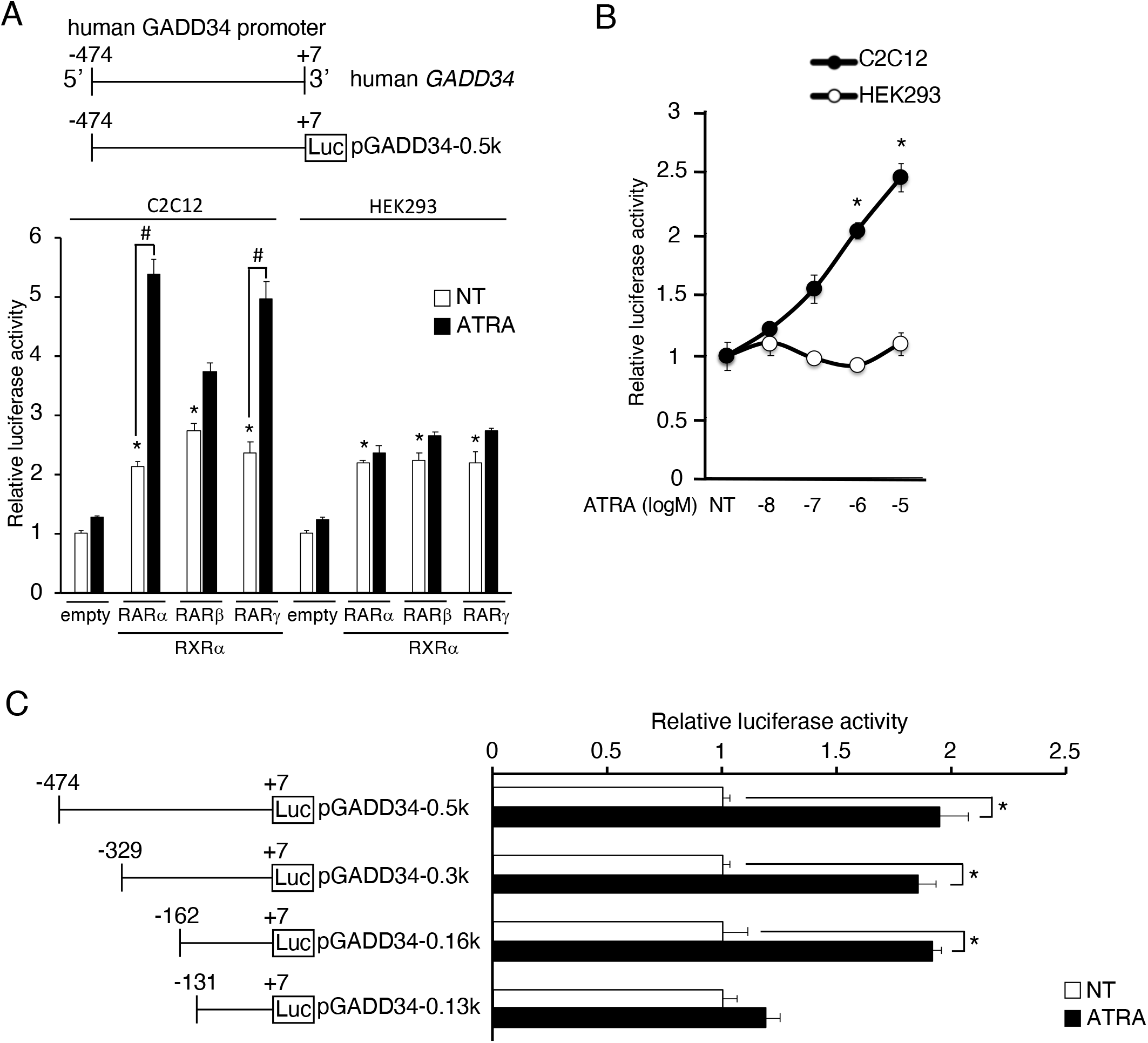
Activation of human *GADD34* gene promoter by ATRA and its receptors in C2C12 cells. A A schematic illustration of the human *GADD34* gene promoter in the upper panels. pGADD34-0.5k and pCMV-β were transfected with pSG5-RAR (α, β, γ) and pSG5-RXRα, or empty vector and incubated in the presence of 1 μM ATRA or DMSO (NT) as a vehicle for 24 h in C2C12 and HEK293 cells (*n* = 3–4). B C2C12 (black circles) and HEK293 (white circle) cells were transfected with pGADD34-0.5k, pSG5-RARα, and pSG5-RXRα, and treated with the indicated concentrations of ATRA for 24 h (*n* = 3–4). C Transcriptional activity of deletion constructs of *GADD34* gene promoters (pGADD34-0.5k, pGADD34-0.3k, pGADD34-0.16k, and pGADD34-0.13k). Deletion constructs are illustrated in the panels on the left. C2C12 cells were transfected with the indicated *GADD34* gene reporter constructs, pSG5-RARα and pSG5-RXRα and incubated in the presence of 1 μM ATRA or DMSO (NT) as a vehicle for 24 h (*n* = 3–4). Data information: In (A), data are presented as the mean ± SEM. \**p* < 0.05 vs. empty vector. #*p* < 0.05 (one-way ANOVA with a Student-Newman post-hoc test). In (B), data are presented as the mean ± SEM. \**p* < 0.05 vs. NT (two-tailed unpaired Student’s *t*-test). In (C), data are presented as the mean ± SEM. \**p* < 0.05 (two-tailed unpaired Student’s *t*-test). Similar results were obtained from independent experiments.

### Six1 downregulates the transcriptional activity of the *GADD34* gene via MEF3-binding site in C2C12 cells

To explore the molecular mechanism underlying the responsiveness of the *GADD34* gene to ATRA in C2C12 myoblasts, we searched for undifferentiated muscle-specific transcription factor binding sites from -162 to -131 on the human *GADD34* gene promoter with Matlnspector-Genomatix. A search for transcription factor binding motifs with this region suggested five potential binding sites: three MEF3 sites, Pax3, and Sox6 (Fig3 A). Based on this search, we overexpressed Six1, a major transcription factor for the MEF3-binding site, Pax3, and Sox6 with GADD34 gene promoter constructs in C2C12 myoblasts. Although each transcription factor decreased the luciferase activity of pGADD34-0.16k, Six1 did not affect the activity of pGADD34-0.13k, unlike Pax3 and Sox6 (Fig3 B). The overexpression of Six1 reduced the GADD34 mRNA and protein expression levels in C2C12 myoblasts (Fig 3C and D). To investigate which MEF3-binding sites are responsible for the decrease in *GADD34* gene promoter activity induced by Six1, we made luciferase reporter vectors of three mutated MEF3-binding sites in the pGADD34-0.13k, as shown in Fig 3E. These mutation analyses showed that the MEF3-2 sequence in the MEF3-binding sites is essential for the repression of the *GADD34* gene promoter activity by Six1 (Fig 3F). Next, we examined whether Six1 binds to the MEF3-2 sequence on the *GADD34* gene promoter with an EMSA analysis. A radiolabeled oligonucleotide containing a GADD34-MEF3 probe, but not a mutated GADD34-MEF3 probe (GADD34 Mut-M2), detected a band in nuclear extracts prepared from C2C12 cells overexpressing Six1 (Fig 3G and H). Although these complexes were susceptible to competition with unlabeled GADD34-MEF3, an unlabeled mutated oligonucleotide (M2: GADD34-Mut-M2) and consensus C/EBP did not compete with these complexes (Fig 3G). These results suggest that Six1 may be involved in the induction of the *GADD34* gene promoter activity by ATRA.

**Figure 3.**
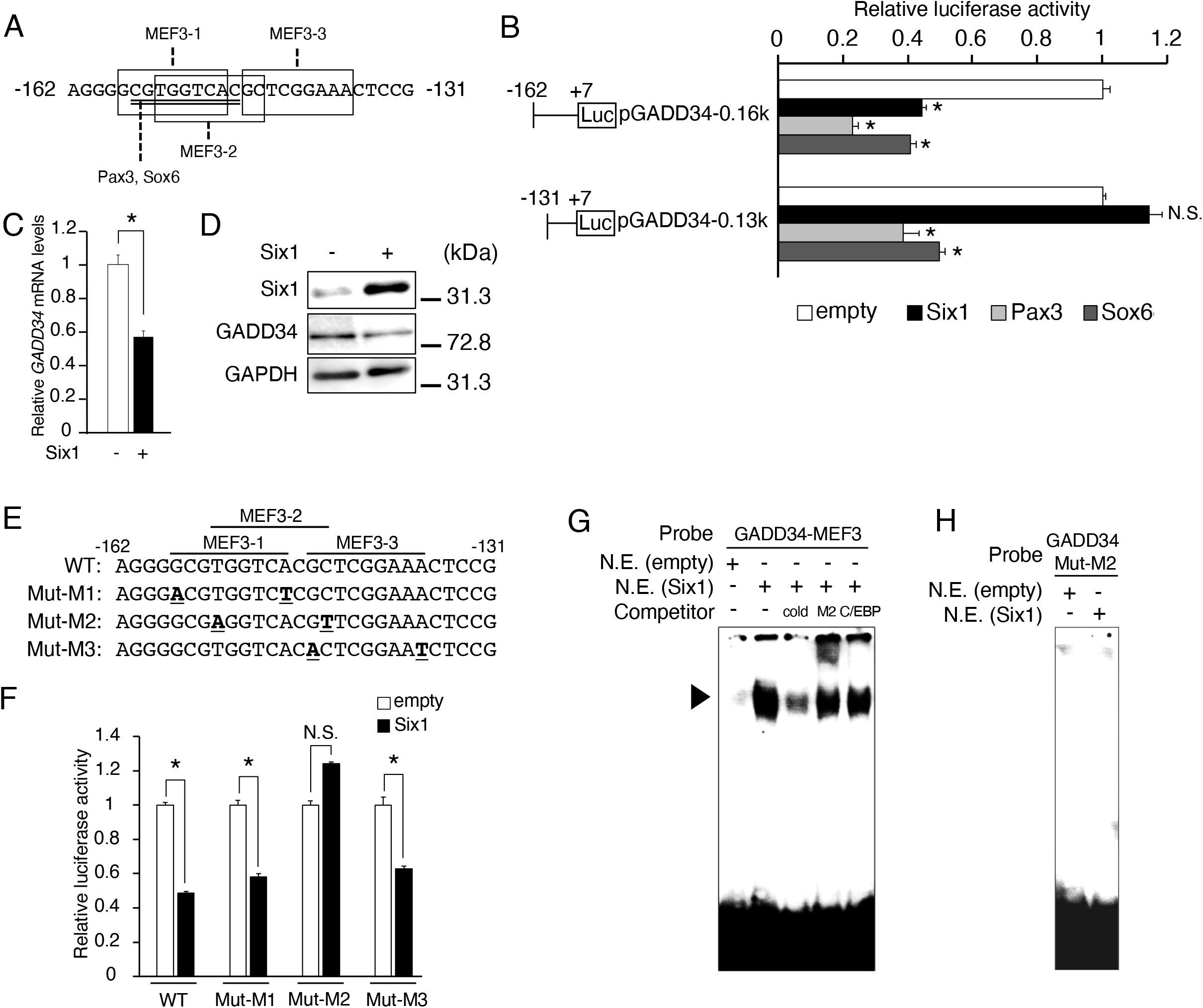
Effects of Six1 on the MEF3 site and the mutation analysis of the *GADD34* gene promoter. A Myoblast-specific transcription factor binding sites (Pax3 and Sox6 sequences: underline, MEF3 sequences: box) in the human *GADD34* gene promoter region. B Each *GADD34* reporter plasmid (pGADD34-0.16k and pGADD34-0.13k) was transfected with each pCR3 plasmid (empty, Six1, Pax3, or Sox6) into C2C12 myoblasts (*n* = 3–4). C The mRNA expression levels of *GADD34* in C2C12 myoblasts transfected with each pCR3 plasmid (empty or Six1) (*n* = 3–4). D Western blotting of Six1 and GADD34 in C2C12 myoblasts transfected with each pCR3 plasmid (empty or Six1). E Three MEF3 sites mutated in the *GADD34* gene reporter region are underlined. Mut-M1, Mut-M2, and Mut-M3 targeted MEF3-1, MEF3-2, and MEF3-3 respectively. F Each *GADD34* reporter plasmid (WT, Mut-M1, Mut-M2, and Mut-M3) was transfected with pcDNA3 empty or Six1 into C2C12 myoblasts (*n* = 3–4). G, H EMSAs using ^32^P-labelled (G) GADD34-MEF3 and (H) mutated GADD34-MEF3-2 (GADD34-Mut-M2) as probes. EMSAs were performed with nuclear extracts (N.E.) from C2C12 myoblasts overexpressing Six1 or empty with the addition of unlabeled competitor oligonucleotides, as indicated. A 100-fold molar excess of each competitor was used. The location of the DNA-protein complex band is indicated by an arrowhead. cold, WT GADD34-Six1; M2, GADD34-Mut-M2; C/EBP, C/EBP-binding sequence. Data information: In (B), data are presented as the mean ± SEM. \**p* < 0.05 vs. empty vector (one-way ANOVA with a Student-Newman post-hoc test). In (C, F), data are presented as the mean ± SEM. \**p* < 0.05 (two-tailed unpaired Student’s *t*-test). N.S. = not significant. Similar results were obtained from independent experiments. Source data for this figure are available online.

### ATRA upregulates the human *GADD34* gene expression through the reduction of the Six1 expression in C2C12 cells

Next, we investigated the effects of ATRA on the Six1 expression in several tissues of mice. The *Six1* mRNA levels were undetectable in the kidney, liver, and EA. ATRA treatment suppressed the mRNA expression of *Six1* (which is highly expressed only in the GM of mice, as well as the protein expression of Six1 (Fig 4A and B). The mRNA expression levels of *Pax3* and *Sox6*, which affect *GADD34* gene promoter activity, were not changed in the GM of ATRA-treated mice in comparison to mice treated with DMSO (Appendix Fig S2A). Likewise, ATRA treatment downregulated the of Six1 mRNA and protein expression in C2C12 myoblasts, but not myotubes (Fig 4C and D). We also examined the time-dependent effects of ATRA on the mRNA expression of *Six1* and *GADD34* for up to 24 h in C2C12 myoblasts. In comparison to DMSO, ATRA transiently decreased the expression of *Six1* mRNA only after 24 h of treatment. It also increased the expression of *GADD34* mRNA; this was observed at both 3 h and 24 h (Fig 4E). The mutated promoter constructs shown in Fig 3E were transfected in C2C12 myoblasts to examine the effects of ATRA on MEF3-binding sites on the *GADD34* gene promoter. These mutation analyses suggested that the binding of Six1 to the MEF3-2 sequence is essential for the activation of the *GADD34* gene promoter by ATRA (Fig 4F). An EMSA assay showed that the band detected by a contact with a radiolabeled oligonucleotide containing a GADD34-MEF3 probe and nuclear extracts prepared from C2C12 cells treated with ATRA was decreased in comparison to treatment with DMSO (Fig 4G). Moreover, Six1-knockdown increased the GADD34 protein expression and inhibited ATRA-induced GADD34 protein expression in C2C12 myoblasts (Fig 4H). As shown in Appendix Fig S1C, since p38 and ERK are involved in the increase of the *GADD34* gene promoter activity by ATRA, we tested whether these MAPK signals regulate the mRNA expression of *Six1* and *GADD34* in myoblasts. As a result, p38 and ERK inhibitors blocked the decrease of the *Six1* mRNA expression and the increase of the *GADD34* mRNA expression induced by ATRA (Appendix Fig S1D). These results indicated that ATRA increases the GADD34 expression via suppression of the binding of Six1 with the MEF3-2 sequence on the human *GADD34* gene promoter by decreasing the expression of Six1.

**Figure 4.**
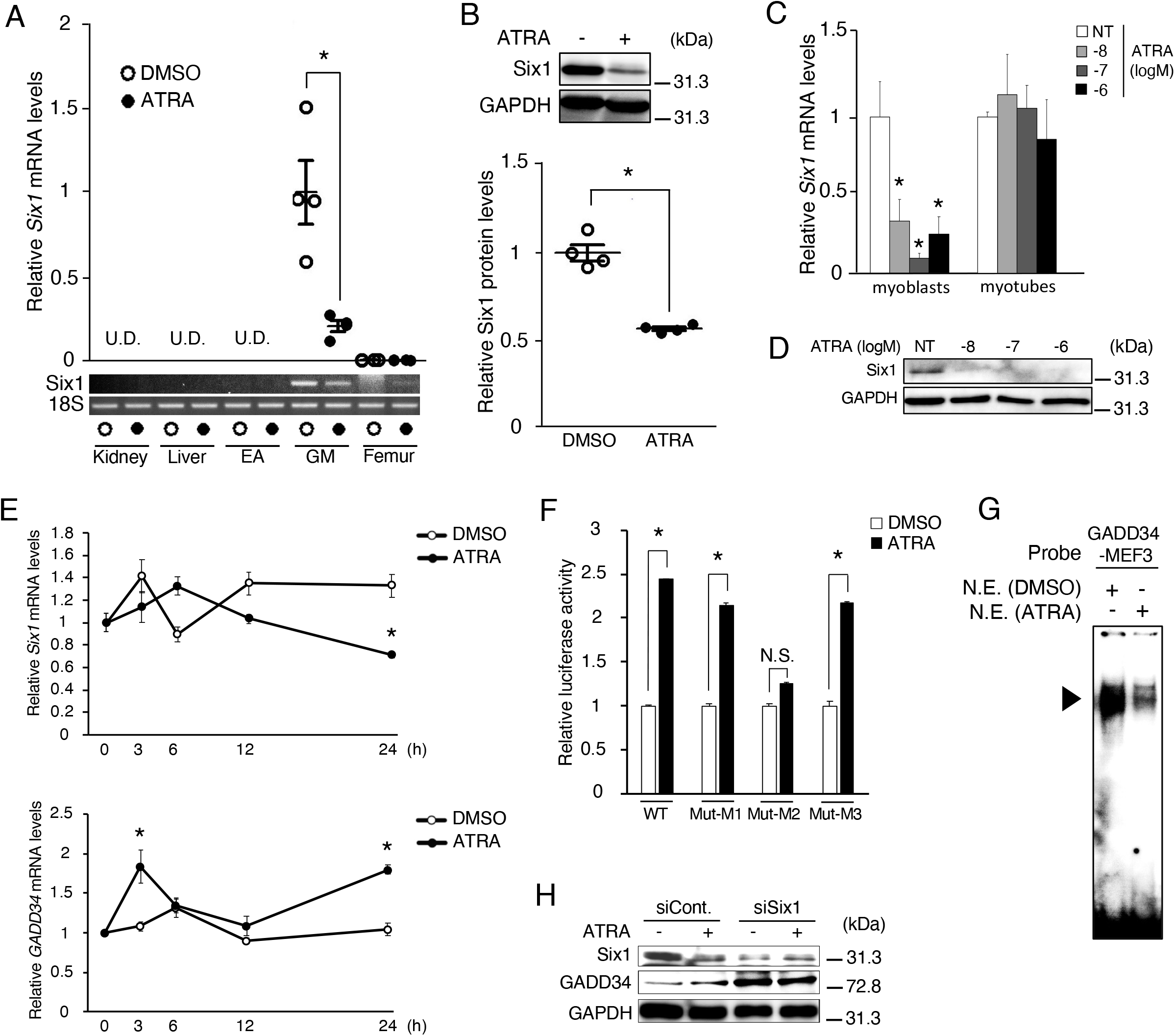
Effects of ATRA treatment on Six1 and GADD34 expression in skeletal muscle. A The mRNA expression levels of *Six1* in kidney, liver, epididymal adipose (EA), gastrocnemius muscle (GM), and femur of ATRA-treated mice (*n* = 4 mice per group). B Western blotting of Six1 in GM of ATRA-treated mice (*n* = 4 mice per group). C The mRNA expression levels of *Six1* in C2C12 cells (myoblasts, myocytes, and myotubes) treated with the indicated concentrations of ATRA (*n* = 3–4). D Western blotting of Six1 in C2C12 myoblasts with the indicated concentrations of ATRA. E The mRNA expression levels of *GADD34* and *Six1* in each C2C12 cells (0, 3, 6, 12, and 24 h) incubated in the presence of 1 μM ATRA or DMSO as a vehicle (*n=*3–4). F Each *GADD34* reporter plasmid (WT, Mut-M1, Mut-M2, and Mut-M3) was transfected and incubated in the presence of 1 μM ATRA or DMSO (NT) as a vehicle for 24 h in C2C12 myoblasts (*n* = 3–4). G EMSAs using ^32^P-labelled GADD34-MEF3 as probes. EMSAs were performed with nuclear extracts (N.E.) from C2C12 myoblasts treated with 1 μM ATRA or DMSO as a vehicle for 24 h. The location of the DNA-protein complex band is indicated by an arrowhead. H Western blotting of Six1 and GADD34 in C2C12 myoblasts transfected with *Six1* siRNA (siSix1) or control (siCont.) and incubated in the presence of 1 μM ATRA or DMSO as a vehicle for 24 h. Data information: In (A, B, F), data are presented as the mean ± SEM. \**p* < 0.05 (two-tailed unpaired Student’s *t*-test). In (C), data are presented as the mean ± SEM. \**p* < 0.05 vs. NT (DMSO) (one-way ANOVA with a Student-Newman post-hoc test). In (E), data are presented as the mean ± SEM. \**p* < 0.05 vs. DMSO (two-tailed unpaired Student’s *t*-test). U.D. = undetectable. N.S. = not significant. Similar results were obtained from independent experiments. Source data for this figure are available online.

### ATRA downregulates the transcriptional activity of the *GADD34* gene via co-repressor TLE3 with Six1 in C2C12 cells

Six1 requires the Eya family as the co-activator for transcriptional activation but requires the TLE family as the co-repressor for transcriptional repression (Jennings & Horowicz, 2008). To define a co-repressor that is responsible for the Six1-mediated reduction of the *GADD34* gene expression, we first analyzed the mRNA expression levels of the TLE family (*TLE1-6*) by qPCR with the absolute standard curve method. Among the TLE family, the *TLE1*, *TLE3*, and *TLE5* genes were expressed in C2C12 myoblasts (Fig 5A), we generated C2C12 cells with the knockdown of these genes using specific siRNA for each of these genes. As shown in Fig 5B, TLE1, TLE3, and TLE5 siRNAs reduced the expression of these genes by 50%, 80%, and 99%, respectively. Although pGADD34-0.16k significantly increased the luciferase activity in response to TLE3-knockdown—but not TLE1-and TLE5-knockdown—pGADD34-0.13k exhibited no increase in response to these knockdowns (Fig 5C). The mutated promoter constructs shown in Fig 3E revealed that the MEF3-2 sequence is essential for the activation of the *GADD34* promoter by TLE3-knockdown (Fig 5D). *TLE3*-knockdown blocked the effects of the luciferase activity of pGADD34-0.16k by the overexpression of Six1 or ATRA treatment (Fig 5E). On the other hand, ATRA did not change the mRNA expression of *TLE1*, *TLE3*, and *TLE5* in C2C12 myoblasts (Appendix Fig S2B). These results indicate that ATRA induces the transcriptional activity of the *GADD34* gene through the inhibition of the collaborative work in TLE3 and Six1 through the decreased expression of Six1.

**Figure 5.**
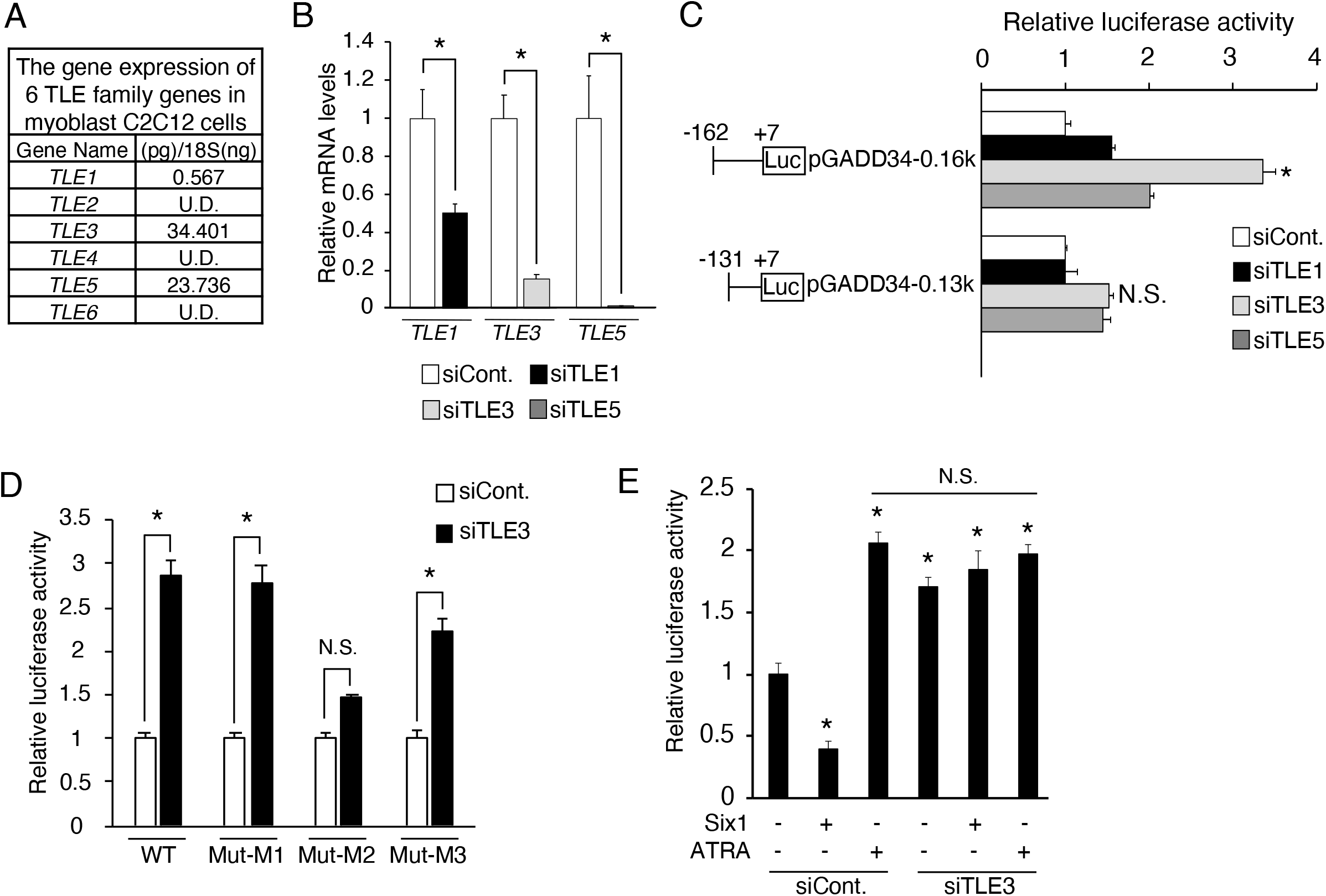
Effects of the TLE family on *GADD34* gene promoter activity in C2C12 myoblasts. A The quantitative expression values of the TLE family (*TLE1–6*) were calculated using the absolute standard curve method of real-time PCR with plasmid templates containing each target gene (*n* = 3). B The mRNA expression levels of *TLE1*, *TLE3*, and *TLE5* in C2C12 myoblasts transfected with each siRNA (siCont., siTLE1, siTLE3, or siTLE5) (*n* = 3–4). C Each *GADD34* reporter plasmid (pGADD34-0.16k and pGADD34-0.13k) was transfected with each siRNA (siCont., siTLE1, siTLE3, or siTLE5) into C2C12 myoblasts (*n* = 3–4). D Each *GADD34* reporter plasmid (WT, Mut-M1, Mut-M2, and Mut-M3) was transfected with *TLE3* siRNA (siTLE3) or control into C2C12 myoblasts (*n* = 3–4). E phGADD34-0.13k was transfected with *TLE3* siRNA (siTLE3) or control and incubated in the presence of 1 μM ATRA or DMSO as a vehicle for 24 h in C2C12 cells (*n* = 3–4). Data information: In (B, D), data are presented as the mean ± SEM. \**p* < 0.05 (two-tailed unpaired Student’s *t*-test). In (C), data are presented as the mean ± SEM. \**p* < 0.05 vs. siCont. (one-way ANOVA with a Student-Newman post-hoc test). In (E), data are presented as the mean ± SEM. \**p* < 0.05 vs. DMSO (NT) (one-way ANOVA with a Student-Newman post-hoc test). U.D. = undetectable. N.S. = not significant.

### ATRA increases the stability of *GADD34* mRNA via TTP in a p38 dependent manner

Thus far, we have demonstrated that ATRA increases the GADD34 expression by transcriptional regulation through the reduced expression of Six1. However, in comparison to vehicle, ATRA transiently increased the *GADD34* mRNA expression after 3 h of treatment, unlike the *Six1* mRNA expression (Fig 4E). Similarly to the mRNA expression, Western blotting revealed that the GADD34 protein expression was increased by ATRA treatment at 3 h in myoblasts (Fig 6A). These results imply the presence of the regulation of GADD34 expression by ATRA, independently of the action of Six1. Next, we tested the stabilization of *GADD34* mRNA at early time points using actinomycin D, a transcriptional inhibitor. As a result, ATRA promoted *GADD34* mRNA stability at 3 h in C2C12 myoblasts that were treated with actinomycin D (Fig 6B). Generally, mRNA stability is regulated through an AU-rich element (ARE: AUUUA motif) in the 3’-untranslated region (3’UTR) of mRNA. To investigate the molecular mechanism underlying the *GADD34* mRNA stabilization in response to ATRA via the 3’UTR of its mRNA, we created a pGL3-GADD34-3’UTR construct (GADD34-3’UTR) by replacing the 3’UTR of the pGL3-basic with the 3’UTR of the *GADD34* gene, which contained two AREs, as illustrated in Fig 6C. ATRA dose-dependently stimulated the luciferase activity of the GADD34-3’UTR, but not pGL3-basic empty (control), in C2C12 myoblasts (Fig 6C).

**Figure 6.**
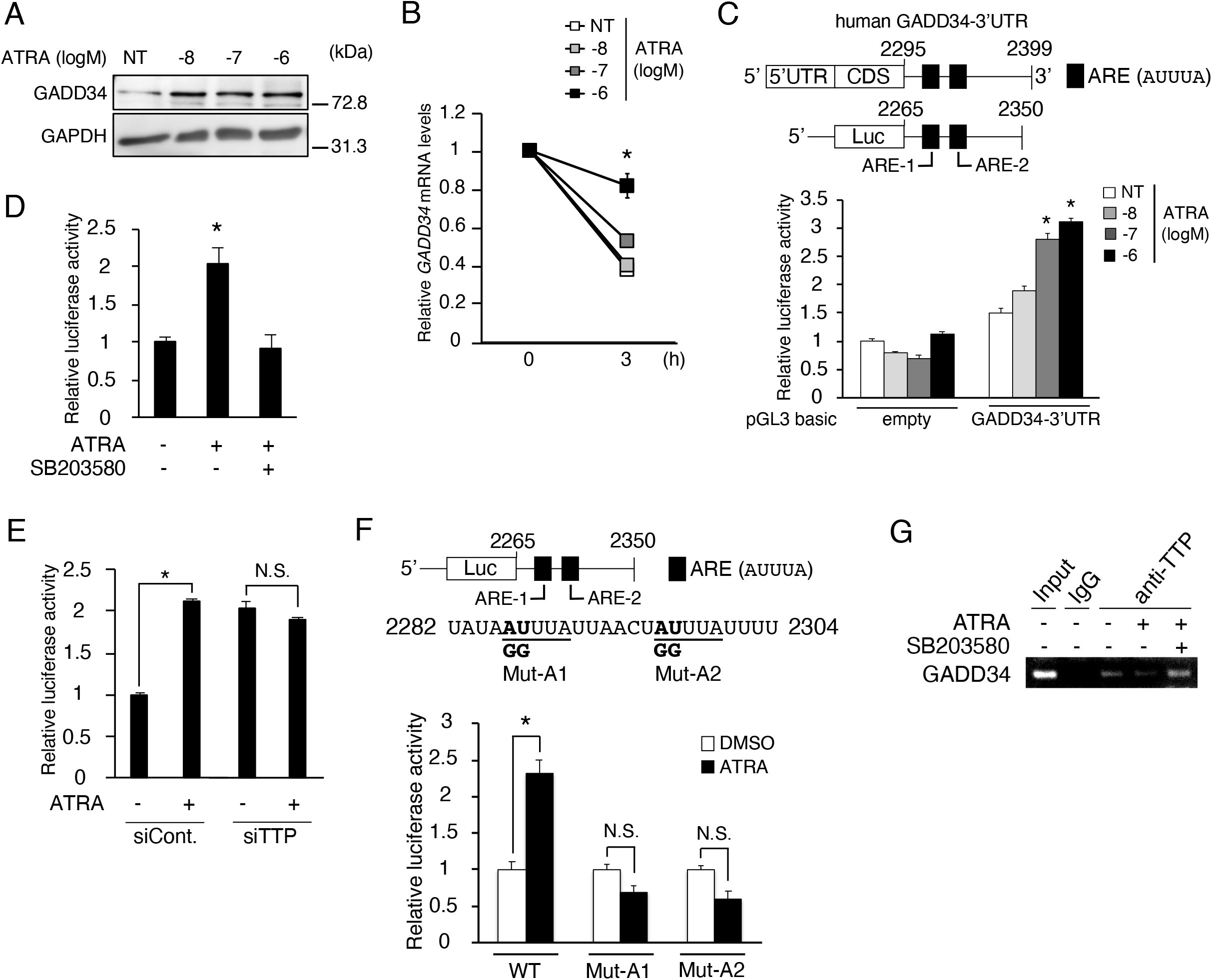
Effects of ATRA on *GADD34* mRNA stability in C2C12 myoblasts. A Western blotting of GADD34 in C2C12 myoblasts incubated in the presence of 1 μM ATRA or DMSO (NT) as a vehicle for 3 h (*n* = 3–4). B The mRNA expression levels of *GADD34* in C2C12 myoblasts treated with the indicated concentrations of ATRA and 2.5 μg/ml Actinomycin D (*n* = 3–4). C A schematic illustration of human *GADD34* mRNA sequence in the upper panels. pGL3-basic plasmid empty (control) or GADD34-3’UTR was transfected and incubated with the indicated concentrations of ATRA for 3 h in C2C12 myoblasts (*n* = 3–4). D, E Human *GADD34* mRNA 3’UTR reporter plasmid was transfected (E) with *TTP* siRNA (siTTP) or control (siCont.) and incubated in the presence of (E) 1 μM ATRA or DMSO (D) and 10 μM SB203580 (p-p38 MAPK inhibitor) for 24 h (*n* = 3–4). F A schematic illustration of the mutated human *GADD34* mRNA sequence in the upper panels. Mut-A1 and Mut-A2 targeted the binding sites AREs for RNA binding protein respectively. Each human *GADD34* mRNA 3’UTR reporter plasmid (WT, Mut-A1, and Mut-A2) was transfected and incubated in the presence of 1 μM ATRA or DMSO as a vehicle for 24 h (*n* = 3–4). G The mRNA expression levels of *GADD34* by RNA ChIP assay in C2C12 cells treated with 1 μM ATRA and 10 μM SB203580 for 24 h (*n* = 3–4). Data information: In (B, C, D, E), data are presented as the mean ± SEM. \**p* < 0.05 vs. NT (DMSO) (one-way ANOVA with a Student-Newman post-hoc test). In (F), data are presented as the mean ± SEM. \**p* < 0.05 (two-tailed unpaired Student’s *t*-test). N.S. = not significant. Similar results were obtained from independent experiments. Source data for this figure are available online.

Since the mRNA stability requires signal transduction pathways of MAPK, such as p38, ERK, and JNK in lung cancer cells (Bhattacharyya *et al*, 2011), we examined which MAPK pathway is responsible for the stabilization of *GADD34* mRNA by ATRA using p38 and ERK inhibitors. Unlike ERK1/2 inhibitor, p38 inhibitor completely suppressed the ATRA-induced luciferase activity of pGADD34-3’UTR (Fig 6D; Appendix Fig S3A and B). The phosphorylation of p38 inhibits the TTP contact with ARE, resulting in the induction of target mRNA stabilization (Bhattacharyya *et al*, 2011). Although ATRA and TNFα, a p38 phosphorylation inducer, stimulated the luciferase activity of GADD34-3’UTR, the effects of these treatments on its activity in C2C12 cells were canceled by TTP-knockdown using TTP-specific siRNA (Fig 6E; Appendix Fig S3C–E). In addition to TTP, human antigen R (HuR), one of the major RBPs, inhibits mRNA degradation (Stoecklin *et al*, 2004). Unlike TTP-knockdown, HuR-knockdown did not change the luciferase activity of GADD34-3UTR induced by ATRA (Appendix Fig S3F and G). To further test whether the ARE1 and ARE2 sequences are responsible for the regulation of *GADD34* mRNA stability by ATRA, we determined the luciferase activity of the mutated ARE1 (Mut-A1) and mutated ARE2 (Mut-A2) in the GADD34-3’UTR (Fig 6F). These mutation analyses showed that both of AREs are important for the enhancement of the stabilization of the *GADD34* mRNA by ATRA (Fig 6F). Using an RNA ChIP assay, we confirmed that p38 inhibitor treatment recovered the decreased binding between *GADD34* mRNA and TTP that was induced by ATRA (Fig 6G). These results suggest that ATRA suppress TTP-induced *GADD34* mRNA degradation through the binding of ARE on the 3’UTR of human *GADD34* mRNA in a p38-dependent manner.

### GADD34 decreases the expression of MYHC2a and changes the muscle fiber type

Finally, C2C12 myoblasts transfected with human GADD34 expression vector or empty (control) at Day 0 were collected at Days 0, 1, 2, and 4 to examine the effects of GADD34 on the C2C12 phenotype (Fig 7A). Unexpectedly, the overexpression of GADD34 did not affect the differentiation speed, thickness of myotubes, the mRNA expression of major muscle differentiation marker genes (*Pax7*, *Myf5*, *MyoD*, and *Myogenin*) or muscle-specific E3 ubiquitin ligase (MuRF1 and Atrogin1), or the synthesis of proteins in C2C12 cells (Fig 7B–D; Appendix Fig S4A–C). Although the GADD34 overexpression did not change the total MYHC protein expression in C2C12 cells from Day 0 to 4, the protein expression of MYHC1 (type 1 slow fibers) was slightly increased in C2C12 cells that overexpressed GADD34 on Day 4. In contrast, the overexpressed GADD34 slightly decreased the protein expression of MYHC2 (type 2 fast fibers) and MYHC2 isoforms, such as MYHC2a and MYHC2x, but not MYHC2b, in C2C12 cells on Day 4 (Fig 7E). In addition, the overexpression of GADD34 downregulated the mRNA expression of *MYH2* and *MYH1*, which respectively encode MYHC2a and MYHC2x proteins (Fig 7F). These results suggest that GADD34 regulates muscle fiber types, such as MYHC2a and MYHC2x, by transcriptional control.

**Figure 7.**
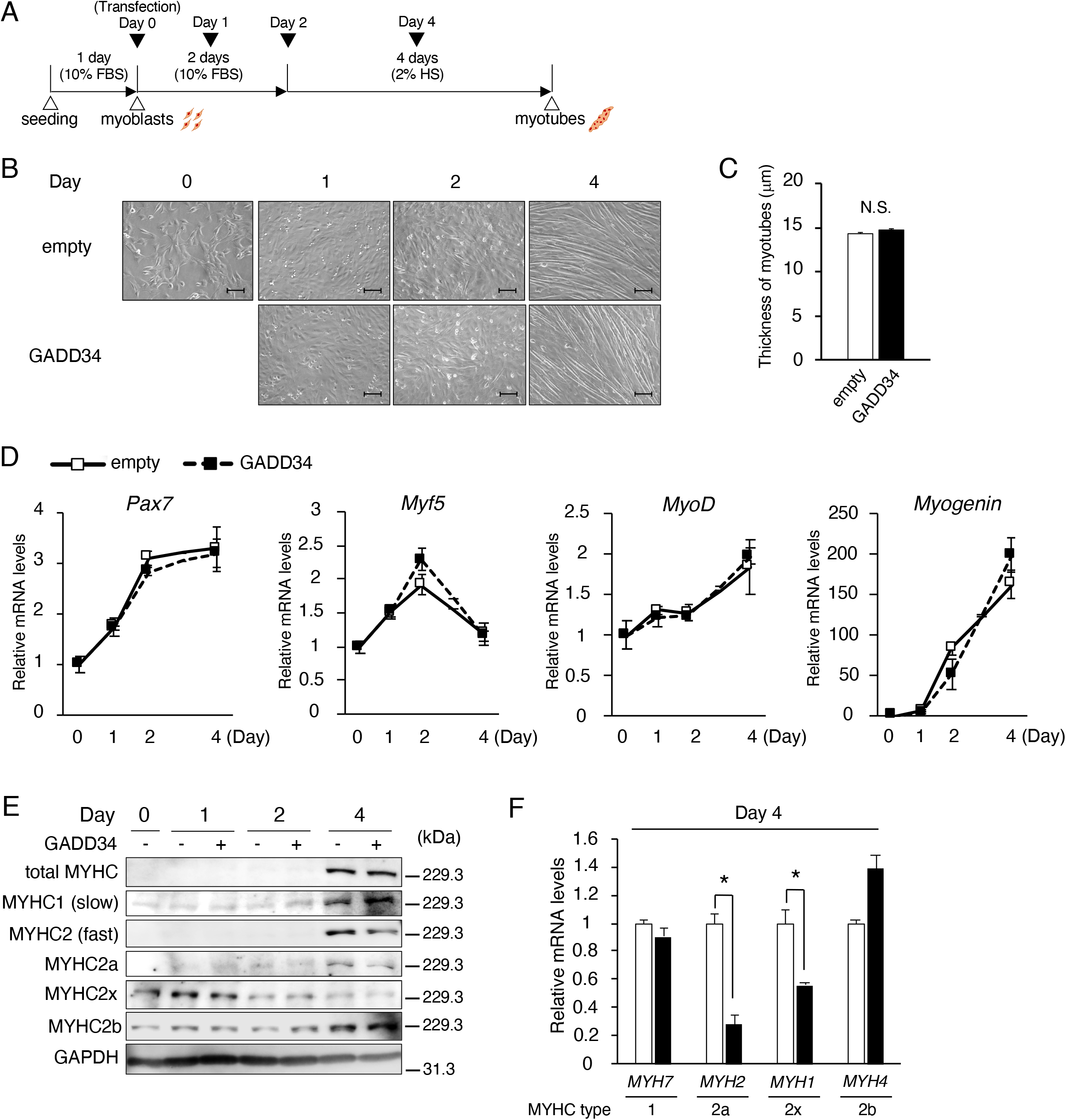
Effects of the overexpression of GADD34 on the muscle fibertype in C2C12 myotubes. A A schematic illustration showing the experimental timeline in C2C12. B Representative images of C2C12 treated as indicated times (Days 0, 1, 2, and 4). Scale bars = 100 μm. C The measurement of diameters of C2C12 myotubes at Day 4, as described in MATERIALS and METHODS (*n* = 100). D The mRNA expression levels of *Pax7, Myf5, MyoD*, and *Myogenin* in C2C12 cells on each day (Days 0, 1, 2, and 4) (*n* = 3–4). E Western blotting of total MYHC, MYHC1 (Slow), MYHC2 (fast), MYHC2a, MYHC2x, and MYHC2b in C2C12 cells on each day (Days 0, 1, 2, and 4). F The mRNA expression levels of *MYH7*, *MYH2*, *MYH1*, and *MYH4* in Day 4 C2C12 cells (*n* = 3–4). Data information: In (C, D, F), data are presented as the mean ± SEM. \**p* < 0.05 (two-tailed unpaired Student’s *t*-test). N.S. = not significant. Source data for this figure are available online.

## Discussion

In the present study, we showed the molecular mechanisms by which ATRA increases the GADD34 expression independently of ATF4 (known to be a potent inducer of GADD34) in myoblasts. Likewise, although a recent study reported that GADD34 levels were decreased in satellite cells collected from injured muscle with increased ATF4 (Xiong *et al*, 2017), the mechanism of this paradoxical result is unknown. Since muscle atrophy can occur with differential sensitivity due to selective skeletal muscle fiber subtypes in various pathological conditions, muscle fiber type change is suggested to be an important factor for muscle wasting (Wanga and Pessinn 2013). Muscle atrophy is a frequent complication in CKD patients (Carrero *et al*, 2013). Interestingly, the plasma levels of vitamin A (retinol or ATRA) increase in CKD patients (Gueguen *et al*, 2005; Jing *et al*, 2016). It has been reported that the percentage of type 1 slow fiber in type 2 fast fiber increases in CKD mice through a decrease in the expression of MYHC2a (Tamaki *et al*, 2014). However, the molecular mechanisms underlying the muscle fiber type change in CKD are largely unknown. In the present study, we revealed that the overexpression of GADD34 increases the MYHC1 protein expression, which is expressed in type 1 slow fibers, and decreases the MYHC2 protein expression, which is expressed in type 2 fast fibers, in myotubes. Based on these results and reports, our findings may provide clues in relation to the unknown mechanisms of pathogenesis, such as muscle atrophy, in CKD patients.

Among the Six family of homeobox (Six1/Six2, Six3/Six6, and Six4/Six5), Six1, Six2, and Six4 are expressed in myoblasts (Kumar, 2009; Grand *et al*, 2012). MyoD reprogramming ability (whereby the MyoD expression turns extra muscle cells into muscle) is impaired in mouse embryonic fibroblasts in Six1/4 double mutant mice because Six1 and MyoD interact with the Myogenin gene promoter to differentiate muscle (Santolini *et al*, 2016). On the other hand, TLE3 downregulates myogenic differentiation via the suppression of MyoD activity (Kokabu *et al*, 2017). Six1 requires the Eya family as the co-activator for transcriptional activation but the TLE family as the co-repressor for transcriptional suppression. In the present study, we suggested that— among the TLE family—*TLE3* is the most abundant gene in myoblasts and is an important co-repressor of Six1 for the regulation of *GADD34* gene transcriptional activity, which consequently changes the ratio of muscle fiber type. In addition, we confirmed that other undifferentiated muscle-specific transcriptional factors, Pax3 and Sox6, also suppress the *GADD34* gene promoter activity. These findings suggest that the expression of GADD34 may constantly be kept low by not only Six1-TLE3 but also by undifferentiated muscle-specific transcriptional factors. However, the reason why the expression of GADD34 is constantly low in myoblasts is unknown.

The expression of genes is determined by transcriptional regulation and post-transcriptional regulation via mRNA stability. The 3’UTR works as one of the regulatory components of mRNA degradation by promoting or inhibiting the deadenylation via interaction with RBP (Mayr, 2019). Although the classic target mRNAs that interact with RBPs for mRNA degradation are associated with inflammatory-related genes, a recent study revealed that other target mRNAs are related to cancer, apoptosis, and other conditions (Kaempfer, 2003; Schuster & Hsieh, 2019; Lal *et al*, 2005). However, UPR-related mRNA has not previously been reported as an ARE-dependent target for stabilization. TTP and HuR, the most famous RBP interacted with ARE (5’-AUUUA-3’) on the 3’UTR, to induce mRNA degradation and stabilization, respectively. However, unlike TTP, HuR did not regulate *GADD34* mRNA stability in the present study. The distinction between HuR and TTP binding has been reported to involve subtle content features: TTP: 5’-AUA[A/U/G/C][A/U]U[A/U]-3’ and HuR: 5’-[A/C/U]UUUU[U/A/C][U/A]-3’ (Bhandare *et al*, 2017). In other words, HuR strongly prefers U-rich sequences, whereas TTP prefers AU-rich with increasing A content. This may explain why HuR did not interact with the ARE sequences of *GADD34* mRNA in the present study. ATRA activates the non-genomic p38 MAPK via extracellular RARα and RARγ (Tanoury *et al*, 2013). Furthermore, phosphorylated p38 inhibits the TTP interaction with ARE on target mRNA (Bhattacharyya *et al*, 2011). Our results indicated that ATRA stabilizes *GADD34* mRNA by inhibiting the interaction between its mRNA and TTP via p38 MAPK. Although it has been reported that ATRA regulates the stabilization of mRNA via interaction between apo-cellular retinoic acid-binding protein 2 (apo-CRABP2) and HuR (Vreeland *et al*, 2014), this is the first study to report that ATRA-dependent MAPK activation increases— via ARE—the mRNA stability of the 3’UTR of the target mRNA. Because p38 MAPK signaling regulates a large number of cellular processes, the mechanism underlying mRNA stabilization by ATRA may also be involved in many mRNAs other than *GADD34* mRNA.

Although the Six genes, especially Six1, are expressed in multiple organs during mammalian development, their expression in global tissues decreases as the individual grows to adulthood (Kumar, 2009). Six1 also controls muscle physiology not only in embryogenesis but also in the adult (Maire *et al*, 2020). In soleus, Six1 deficiency reduced MYHC2a fiber in 3-week-old C57BL6 mice and caused the complete disappearance of the expression of MYHC2a fiber in 12-week-old C57BL6 mice (Sakakibara *et al*, 2016). These results imply that Six1 regulates fast-fiber type acquisition and maintenance in adult mice. Although the effects of GADD34 on skeletal muscle *in vivo* are unknown, our study revealed that GADD34, which is downregulated by Six1, suppressed the MYHC2a expression in myoblasts. Thus, GADD34 may mediate the regulation of the expression of MYHC2a by Six1 in myotubes. Aside from this, GADD34 induces cellular senescence via the regulation of the p21 expression in MEF and 7EJ-Ras cells (Minami *et al*, 2007). Cellular senescence, a permanent state of replicative arrest in otherwise proliferating cells, is a hallmark of aging and has been linked to aging-related diseases (Childs *et al*, 2015). Since the plasma levels of vitamin A (retinol or ATRA) increase with aging (Gueguen *et al*, 2005), the increase of ATRA that occurs with aging may accelerate cellular senescence in the muscle via the induction of GADD34, resulting in muscle atrophy in elderly patients. It has been reported that aged mice exhibit an increase of slow muscle fibers but a decrease of fast muscle fibers (Shang *et al*, 2020). These results are in line with the effects of GADD34 on myotubes that were observed in our study.

In conclusion, as shown in Fig 8, our findings revealed that the upregulation of the *GADD34* gene expression by ATRA contributes to muscle fiber type change in myoblasts. Furthermore, ATRA and its receptors can increase the transcriptional activity of the *GADD34* gene by blocking the interaction of Six1-TLE3 with the *GADD34* gene promoter and stabilize *GADD34* mRNA by inhibiting the interaction of p38-TTP with the 3’UTR of *GADD34* mRNA.

**Figure 8.**
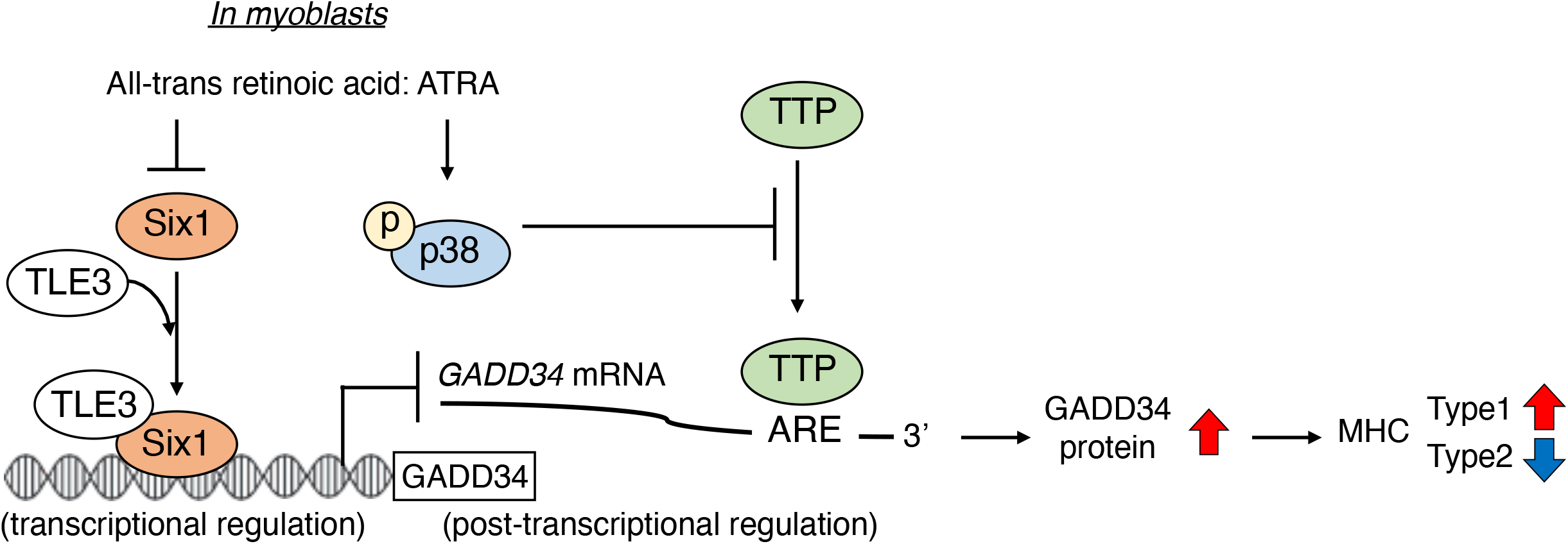
Schematic illustration of muscle specific GADD34 induction through ATRA dependent transcriptional regulation and post-transcriptional regulation. ATRA decreases the expression of Six1, which decreases the transcriptional activity of *GADD34* with TLE3 in myoblasts. ATRA also increases *GADD34* mRNA stability via p38 phosphorylation, which leads to the blocking of interaction between TTP and ARE on *GADD34* mRNA. Increased GADD34 proteins change the type of myosin heavy chain in myotubes.

## Materials and Methods

### Chemicals and reagents

ATRA, DMSO, FR180204, anti-GAPDH antibody, anti-Six1 antibody, goat anti-mouse IgG (H + L)-HRP conjugate, goat anti-rabbit IgG (H + L)-HRP conjugate, MISSION siRNA oligos were purchased from Sigma-Aldrich (St. Louis, MO, USA). Buprenorphine hydrochloride was purchased from Otsuka Pharmaceutical Co., Ltd. (Tokyo, Japan). Pentobarbital sodium salt was purchased from Tokyo Kasei Co., Ltd. (Tokyo, Japan). Anti-phosphorylated eIF2α (p-eIF2α) (#9721, Ser51), anti-eIF2α (#9722), anti-ATF4 (#11815), anti-CHOP (#2895), anti-TTP (#71632), anti-phosphorylated ERK (p-ERK) (#9101, Thr202/Try204), and anti-ERK (#9102) antibody were purchased from Cell Signaling Technology (Danvers, MA, USA). Anti-phosphorylated p38 (p-p38) (sc-166182, Tyr182), anti-p-38 (sc-7972), and anti-HuR (sc-5261) antibody, and C/EBP consensus oligonucleotide (cebp; catalog number sc-2525) were purchased from Santa Cruz Biotechnology (Santa Cruz, CA, USA). Anti-GADD34 antibody (10449-1-AP) was purchased from Proteintech (Rosemont, IL, USA). Anti-MYHC1 (BA-D5), anti-MYHC2 (F59), anti-MYHC2a (SC-71), anti-MYHC2x (6H1), and anti-MYHC2b (BF-F3) antibody were purchased from Developmental Studies Hybridoma Bank (Iowa city, IA, USA). Anti-puromycin antibody was purchased from Cosmo Bio (Tokyo, Japan). [γ-32P] ATP was purchased from ICN (Costa Mesa, CA, USA). T4 polynucleotide kinase and TransIT®-LT1 Reagent were purchased from TAKARA (Shiga, Japan). SB203580 was purchased from Adipogen Life Science (San Diego, CA, USA). Human recombinant TNFα was purchased from Invitrogen (Carlsbad, CA, USA). Puromycin was purchased from Funakoshi (Tokyo, Japan). Chemi-Lumi One Super was purchased from Nacalai Tesque (Kyoto, Japan). QIAzol^®^ Lysis Reagent was purchased from QIAGEN (Venlo, Nederland). TOYOBO KOD one™ PCR Master Mix -Blue-was purchased from TOYOBO (Osaka, Japan). RNA ChIP-IT^®^ was purchased from Active Motif (Tokyo, Japan).

### Animal experiments

The animal work took place in Division for Animal Research and Genetic Engineering Support Center for Advanced Medical Sciences, Institute of Biomedical Sciences, Tokushima University Graduate School. Eight-week-old male C57BL/6J mice were purchased from Japan SLC (Shizuoka, Japan) and individually caged in a climate-controlled room (22 ± 2°C) with a 12-h light-dark cycle. These mice were randomly divided into two groups and a total of 1 mg/kg body weight of ATRA or 1% DMSO (control) prepared in sterile saline was intraperitoneally administered, then the mice were sacrificed 24 h later. Before sacrifice, mice were anesthetized with a total of 0.1 mg/kg body weight of buprenorphine hydrochloride and a total of 50 mg/kg body weight of pentobarbital sodium salt, and tissues were removed. The present study was approved by the Animal Experimentation Committee of Tokushima University School of Medicine (animal ethical clearance No. T30-66) and was carried out in accordance with guidelines for the Animal Care and use Committee of Tokushima University School of Medicine.

### Cell culture

C2C12 myoblast cells were cultured as described previously (Niida *et al*, 2020). Briefly, C2C12 myoblast cells were cultured in DMEM (Sigma, St. Louis, MO) containing 10% FBS (Sigma), 100 units/ml penicillin and 100 µg/ml streptomycin at 37°C with 5% CO_2_. At 100% confluence, C2C12 myoblast cells were fused by shifting the medium to DMEM containing 2% horse serum (HS; Moregate Biotech, Bulimba, Australia). These cells were maintained in 2% HS/DMEM (differentiation medium) for 4 days prior to experiments. HEK293 cells were cultured in DMEM containing 10% FBS, 100 units/ml penicillin and 100 µg/ml streptomycin at 37°C with 5% CO_2_.

### Western blotting

Protein samples were heated to 95°C for 5 min in sample buffer in the presence of 5% 2-mercaptoethanol and subjected to SDS–PAGE. The separated proteins were transferred by electrophoresis to polyvinylidene difluoride transfer membranes (Immobilon-P, Millipore, MA, USA). The membranes were treated with diluted affinity-purified anti-p-eIF2α (p-eIF2α) (1:1000), anti-eIF2α (1:1000), anti-ATF4 (1:3000), anti-CHOP (1:1000), anti-GADD34 (1:1000), anti-Six1 (1:1000), anti-TTP (1:1000), anti-HuR (1:1000), anti-p-p38 (p-p38) (1:1000), anti-p38 (1:1000), anti-p-ERK (p-ERK) (1:1000), anti-ERK (1:1000), anti-MYHC1 (1:1000), anti-MYHC2 (1:1000), anti-MYHC2a (1:1000), anti-MYHC2x (1:500), anti-MYHC2b (1:500), and anti-puromycin (1:2000) antibody. Mouse anti-GAPDH monoclonal antibody was used as an internal control. Goat anti-mouse IgG (H + L)-HRP conjugate (1:3000) and Goat anti-rabbit IgG (H + L)-HRP conjugate (1:3000) was utilized as the secondary antibody, and signals were detected using the Chemi-Lumi One Super.

### Quantitative PCR

Total RNA was isolated from kidney, liver, EA, GM, femur, C2C12 myoblasts, C2C12 myotubes, and HEK293 cells using an QIAzol^®^ Lysis Reagent. Real-time quantitative PCR assays were performed using an Applied Biosystems StepOne qPCR instrument. In brief, the cDNA was synthesized from 1 μg of total RNA using a reverse transcriptase kit (Invitrogen, Carlsbad, CA) with an oligo-dT primer. After cDNA synthesis, quantitative real-time PCR was performed in 5 μl of SYBR Green PCR master mix using a real-time PCR system (Applied Biosystems). Amplification products were then analyzed by a melting curve, which confirmed the presence of a single PCR product in all reactions (apart from negative controls). The quantification of given genes was expressed as the mRNA level normalized to a ribosomal 18S house keeping gene using the ΔΔCt method. Quantitative expression values were calculated from an absolute standard curve method using the plasmid template for each target gene. The primer sequences used for real-time PCR analysis are shown in Appendix Table S1.

### Reporter plasmid construction

The promoter fragment of luciferase reporter plasmid pGADD34-0.5k generated by Eurofins Japan (Tokyo, Japan) was subcloned into a pGL-3 vector (Promega, Madison, WI, USA) by restriction enzyme cutting site KpnI/HindIII. Luciferase reporter plasmids GADD34-3’UTR was constructed by PCR amplification of human genomic cDNA as a template using gene-specific primers (Appendix Table S2). These products were subcloned into a pGL-3 vector. Deleted reporter plasmid pGADD34-0.3k was cloned by the digestion of pGADD34-0.5k using DpnI/HindIII. Deleted reporter plasmids pGADD34-0.16k, pGADD34-0.13k, and mutated reporter plasmids pGADD34-0.16k-Mut-MEF3-1 (Mut-M1), pGADD34-0.16k-Mut-MEF3-2 (Mut-M2), pGADD34-0.16k-Mut-MEF3-3 (Mut-M3), GADD34-3’UTR-Mut-ARE-1 (Mut-A1), GADD34-3’UTR-Mut-ARE-2 (Mut-A2) were constructed with TOYOBO KOD one^TM^ PCR Master Mix -Blue-using the oligonucleotides shown in Appendix Table S2. The β-galactosidase expression vector pCMV-β (CLONTECH, Palo Alto, CA, USA) was used as an internal control. Each plasmid was purified with a FavorPrep™ Plasmid DNA Extraction Midi Kit (Favorgen, Ping-Tung, Taiwan).

### Transfection and luciferase assay

Mouse Six1, Pax3, and Sox6 expression vector (pCR3-Six1, pCR3-Pax3, and pCR3-Sox6) were kindly provided by Dr. P. Maire (Santolini *et al*, 2016). Human GADD34 expression vector (pRP-Neo-CMV-hPPP1R15A) was designed by Vector Builder. Cells were transfected by TransIT^®^-LT1 Reagent, then treated with several concentrations of ATRA, SB203580, FR180204, TNFα, or DMSO as vehicle control for an additional 16 h. A luciferase assay was performed as described previously (Masuda *et al*, 2010).

### RNAi experiments

C2C12 myoblasts were transfected with siRNA directed against ATF4 (SASI_Mm02_00316863), Six1 (SASI_Mm01_00198104), TLE1 (SASI_Mm01_00069933), TLE3 (SASI_Mm02_00300046), TLE5 (SASI_Mm01_00139428), TTP (SASI_Mm01_00178605), HuR (SASI_Mm02_00318722), or negative control (SIC001) using TransIT®-LT1 Reagent, according to the manufacturer’s instructions.

### EMSA

EMSA was performed as described previously (Masuda *et al*, 2020). Double-stranded nucleotides for GADD34-MEF3 and GADD34-MEF3-2-Mutant (GADD34-Mut-M2) were synthesized (Appendix Table S3). Purified DNA fragments were radiolabeled with [γ-32P] ATP (110 TBq/mmol) using T4 polynucleotide kinase. Nuclear extracts (pCR3-Six1) were prepared as described previously (Masuda *et al*, 2010). Briefly, the C2C12 myoblast cells were cultured in 10-cm dishes to 60% confluence and transfected with pCR3-Six1 or treated with 1 μM ATRA. Prepared nuclear extracts (15 μg) were incubated with the radiolabeled probe in binding buffer [10 mM (Tris–HCl), pH7.5, 1 mM DTT, 1 mM EDTA, 10% Glycerol, 1 mM MgCl2, 0.25 mg/ml bovine serum albumin, 2.5 μg/ml salmon sperm DNA and 2 μg poly(dI-dC)] in a final volume of 20 μl for 30 min at room temperature. The specificity of the binding reaction was determined with a 100-fold molar excess of the indicated cold competitor oligonucleotide. The reaction mixture was then subjected to electrophoresis on a 5% polyacrylamide gel with 0.25×TBE running buffer for 2 h at 150 V. The gel was dried and analyzed with an image scanner (FLA-9000 Starion, Tokyo, Japan).

### RNA ChIP assay

Using the RNA ChIP-IT^®^, an RNA ChIP assay was performed according to the manufacturer’s instructions. Briefly, the C2C12 myoblast cells were cultured to 80% confluence in 10-cm dishes and treated with 1 μM ATRA. The C2C12 cells were collected and lysed in lysis buffer. The cell extract was prepared and incubated with RNA ChIP buffer pre-conjugated with TTP antibodies or control mouse IgG at 4°C for 16 h. The complexes were treated with Proteinase K for 1 h at 45°C and 1.5 h at 65°C. Immunoprecipitated RNA in the precipitates was purified using QIAzol^®^ Lysis reagent and analyzed for GADD34 by RT-qPCR.

### Measurement of C2C12 diameters

The myotube diameters were determined as previously reported (Niida *et al*, 2020). Under a fluorescence microscope (BIOREVO BZ-9000, Keyence), three photographs per cell-culture well were obtained in a high-power field. We measured the diameter at the middle portion of the myotube with the built-in BZ-II analyzer software program. We measured the diameters of 100 myotubes/group.

### Surface Sensing of Translation (SUnSET) assay in C2C12 cells

C2C12 was incubated with or without 1 μM puromycin for 30 min before collecting the cells and washed with PBS as previously described (Lim *et al*, 2017). Puromycin-labeled proteins were detected by Western blotting, as shown above.

### Statistical analysis

Data were collected from more than two independent experiments and were reported as the mean and SEM. The statistical analysis for two-group comparison was performed using a two-tailed Student’s *t*-test, or one-way ANOVA with Tukey-Kramer post-hoc test for multi-group comparison. All data analyses were performed using the GraphPad Prism 5 software program (GraphPad Software, San Diego, CA, USA). *P* values of < 0.05 were considered to indicate statistical significance.

## Reference

Alter J, Bengal E (2011) Stress-induced C/EBP homology protein (CHOP) represses MyoD transcription to delay myoblast differentiation. PLoS One 6: e29498

Bhandare S, Goldberg DS, Dowell R (2017) Discriminating between HuR and TTP binding sites using the k-spectrum kernel method. PLos One 12: e0174052

Bhattacharyya S, Gutti U, Mercado J, Moore C, Pollard HB, Biswas R (2011) MAPK signaling pathways regulate IL-8 mRNA stability and IL-8 protein expression in cystic fibrosis lung epithelial cell lines. Am J Physiol Lung Cell Mol Physiol 3000: L81–87

Bouchard F, Paquin J (2013) Differential effects of retinoids and inhibitors of ERK and p38 signaling on adipogenic and myogenic differentiation of P19. stem cells 22: 2003–2016

Carreras-Sureda A, Hetz C (2015) RNA metabolism: putting the brake on the UPR. EMBO Rep 16: 545–546

Carrero JJ, Stenvinkel P, Cuppari L, Ikizler TA, Kalantar-Zadeh K, Kaysen G, Mitch WE, Price SR, Wanner C, Wang AYM et al (2013) Etiology of the protein-energy wasting syndrome in chronic kidney disease: a consensus statement from the International Society of Renal Nutrition and Metabolism (ISRNM). J Ren Nutr 23: 77–90

Chambon P (1996) A decade of molecular biology of retinoic acid receptors. FASEB J 10: 940–954

Chen J, Wang Y, Hamed M, Lacroix N, Li Q (2015) Molecular Basis for the Regulation of Transcriptional Coactivator p300 in Myogenic Differentiation. Sci Rep 5: 13727

Childs BG, Durik M, Baker DJ, Deursen JM (2015) Cellular senescence in aging and age-related disease: from mechanisms to therapy. Nat Med 12:1424–1435

Clagett-Dame M, DeLuca FH (2002) The role of vitamin A in mammalian reproduction and embryonic development. Annu Rev Nutr 22: 347–381

Clavarino G, Cláudio N, Dalet A, Terawaki S, Couderc T, Chasson L, Ceppi M, Schmidt EK, Wenger T, Lecuit M et al (2012) Protein phosphatase 1 subunit Ppp1r15a/GADD34 regulates cytokine production in polyinosinic: polycytidylic acid-stimulated dendritic cells. Proc Natl Acad Sci U S A 109: 3006–3011

Ebert SM, Bullard SA, Basisty N, Marcotte GR, Skopec ZP, Dierdorff JM, Al-Zougbi A, Tomcheck KC, DeLau AD, Rathmacher JA et al (2020) Activating transcription factor 4 (ATF4) promotes skeletal muscle atrophy by forming a heterodimer with the transcriptional regulator C/EBPβ. J Biol Chem 295: 2787–2803

Eochette-Egly C (2015) Retinoic acid signaling and mouse embryonic stem cell differentiation: Cross talk between genomic and non-genomic effects of RA. Biochim Biophys Acta 1851: 66–75

Gallot YS, Bohnert KR, Straughn AR, Xiong G, Hindi SM, Kumar A (2019) PERK regulates skeletal muscle mass and contractile function in adult mice. FASEB J 33: 1946-1962

Goh CW, Lee IC, Sundaram JR, George SM, Yusoff P, Brush MH, Sze NSK, Shenolikar S (2018) Chronic oxidative stress promotes GADD34-mediated phosphorylation of the TAR DNA-binding protein TDP-43, a modification linked to neurodegeneration. J Biol Chem 293: 163–176

Grand FL, Grifone R, Mourikis P, Houborn C, Gigaud C, Pujol J, Maillet M, Pagès G, Rudnicki M, Tajbakhsh S et al (2012) Six1 regulates stem cell repair potential and self-renewal during skeletal muscle regeneration. J Cell Biol 198: 815–832

Gueguen S, Leroy P, Gueguen R, Siest G, Visvikis S, Herbeth B (2005) Genetic and environmental contributions to serum retinol and alpha-tocopherol concentrations: the Stanislas Family Study. Am J Clin Nutr 81: 1034–1044

Harding HP, Novoa I, Zhang Y, Zeng H, Wek R, Schapira M, Ron D (2000) Regulated translation initiation controls stress-induced gene expression in mammalian cells. Mol Cell 6: 1099–1108

Harding HP, Zhang Y, Scheuner D, Chen JJ, Kaufman RJ, Ron D (2009) Ppp1r15 gene knockout reveals an essential role for translation initiation factor 2 alpha (eIF2alpha) dephosphorylation in mammalian development. Proc Natl Acad Sci U S A 106: 1832-1837

Jennings BH, Horowicz DI (2008) The Groucho/TLE/Grg family of transcriptional co-repressors. Genome Biol 9: 205

Jing J, Isoherranen N, Robinson-Cohen C, Petrie I, Kestenbaum BR, Yeung CK (2016) Chronic Kidney Disease Alters Vitamin A Homeostasis via Effects on Hepatic RBP4 Protein Expression and Metabolic Enzymes. Clin Transl Sci 9: 207–215

Kaempfer R (2003) RNA sensors: novel regulators of gene expression. EMBO Rep 4: 1043–1047

Kim J, Wellmann KB, Smith ZK, Johnson BJ (2018) All-trans retinoic acid increases the expression of oxidative myosin heavy chain through the PPARδ pathway in bovine muscle cells derived from satellite cells. J Anim Sci 96: 2763–2776

Khatib T, Marini P, Nunna S, Chisholm DR, Whiting A, Redfern C, Greig IR, McCaffery P (2019) Genomic and non-genomic pathways are both crucial for peak induction of neurite outgrowth by retinoids. Cell Commun Signal 17: 40

Kim I, Xu W, Reed JC (2008) Cell death and endoplasmic reticulum stress: disease relevance and therapeutic opportunities. Nat Rev Drug Discov 7: 1013–1030

Kokabu S, Nakatomi C, Matsubara T, Ono Y, Addison WN, Lowery JW, Urata M, Hudnall AM, Hitomi S, Nakatomi M et al (2017) The transcriptional co-repressor TLE3 regulates myogenic differentiation by repressing the activity of the MyoD transcription factor. J Biol Chem 292: 12885–12894

Krokowski D, Guan BJ, Wu J, Zheng Y, Pattabiraman PP, Jobava R, Gao XH, Di XJ, Snider MD, Mu TW, et al (2017) GADD34 Function in Protein Trafficking Promotes Adaptation to Hyperosmotic Stress in Human Corneal Cells. Cell Rep 21: 2895–2910

Kumar JP (2009) The sine oculis homeobox (SIX) family of transcription factors as regulators of development and disease. Cell Mol Lif Sci 66: 565–583

Lal A, Kawai T, Yang X, Mazan-Mamczarz K, Gorospe M (2005) Antiapoptotic function of RNA-binding protein HuR effected through prothymosin alpha. EMBO J 24: 1852–1862

Lamarche É, Lala-Tabbert N, Gunanayagam A, St-Louis C, Wiper-Bergeron N (2015) Retinoic acid promotes myogenesis in myoblasts by antagonizing transforming growth factor-beta signaling via C/EBPβ. Skelet Muscle 5: 8

Lim JA, Li L, Shirihai OS, Trudeau KM, Puertollano R, Raben N (2017) Modulation of mTOR signaling as a strategy for the treatment of Pompe disease. EMBO Mol Med 9: 353–370

Maire P, Santos MD, Madani R, Sakakibara I, Viaut C, Wurmser M (2020) Myogenesis control by SIX transcriptional complexes. Semin Cell Dev Biol 104: 51–64

Mangelsdorf DJ, Evans RM (1995) The RXR heterodimers and orphan receptors. Cell 83: 841–850

Masciarelli S, Capuano E, Ottone T, Divona M, Panfilis SD, Banella C, Noguera NI, Picardi A, Fontemaggi G, Blandino G et al (2018) Retinoic acid and arsenic trioxide sensitize acute promyelocytic leukemia cells to ER stress. Leukemia 32: 285–294

Masuda M, Yamamoto H, Kozai M, Tanaka S, Ishiguro M, Takei Y, Nakahashi O, Ikeda S, Uebanso T, Taketani Y et al (2010) Regulation of renal sodium-dependent phosphate co-transporter genes (Npt2a and Npt2c) by all-trans-retinoic acid and its receptors. Biochem J 429: 583–592

Masuda M, Miyazaki-Anzai S, Levi M, Ting TC, Miyazaki M (2013) PERK-eIF2a-ATF4-CHOP signaling contributes to TNFa-induced vascular calcification. J Am Heart Assoc 2: e000238

Masuda M, Miyazaki-Anzai S, Keenan AL, Okamura K, Kendrick J, Chonchol M, Offermanns S, Ntambi JM, Kuro-O M, Miyazaki M (2015) Saturated phosphatidic acids mediate saturated fatty acid-induced vascular calcification and lipotoxicity. J Clin Invest 125: 4544-4558

Mayr C (2019) What Are 3’ UTRs Doing? Cold. Spring Harb Perspect Biol 11: a034728

Minami K, Inoue H, Terashita T, Kawakami T, Watanabe R, Haneda M, Isobe K, Okabe H, Chano T (2007) GADD34 induces p21 expression and cellular senescence. Oncol Rep 17: 1481–1485

Moore K, Hollien J (2015) Ire1-mediated decay in mammalian cells relies on mRNA sequence, structure, and translational status. Mol Biol Cell 26: 2873-2884

Nair SS, Das SS, Nair RP, Indira M (2018) Supplementation of all trans retinoic acid ameliorates ethanol-induced endoplasmic reticulum stress. Arch Physiol Biochem 124: 131–138

Niida Y, Masuda M, Adachi Y, Yoshizawa A, Ohminami H, Mori Y, Ohnishi K, Yamanaka-Okumura H, Uchida T, Nikawa T et al (2020) Reduction of stearoyl-CoA desaturase (SCD) contributes muscle atrophy through the excess endoplasmic reticulum stress in chronic kidney disease. J Clin Biochem Nutr 67: 179–187

Nishio N, Isobe K (2015) GADD34-deficient mice develop obesity, nonalcoholic fatty liver disease, hepatic carcinoma and insulin resistance. Sci Rep 5: 13519

Pakos-Zebrucka K, Koryga I, Mnich K, Ljujic M, Samali A, Gorman AM (2016) The integrated stress response. EMBO Rep 17: 1374–1395

Rayavarapu S, Coley W, Nagaraju K (2012) Endoplasmic reticulum stress in skeletal muscle homeostasis and disease. Curries Rheumatol Rep 14: 238–243

Sakakibara I, Wurmser M, Santos MD, Santolini M, Ducommun S, Davaze R, Guernec A, Sakamoto K, Maire P (2016) Six1 homeoprotein drives myofiber type IIA specialization in soleus muscle. Skelet Muscle 6: 30

Santolini M, Sakakibara I, Gauthier M, Ribas-Aulinas F, Takahashi H, Sawasaki T, Mouly V, Concordet JP, Defossez PA, Hakim V et al (2016) MyoD reprogramming requires Six1 and Six4 homeoproteins: genome-wide cis-regulatory module analysis. Nucleic Acids Res 44: 8621–8640

Sartori R, Romanello V, Sandri M (2021) Mechanisms of muscle atrophy and hypertrophy: implications in health and disease. Nat Commun 12: 330

Schuster SL, Hsieh AC (2019) The Untranslated Regions of mRNAs in Cancer. Trends Cancer 4: 245–262

Shang GK, Han L, Wang ZH, Liu YP, Yan SB, Sai WW, Wang D, Li YH, Zhang W, Zhong M (2020) Sarcopenia is attenuated by TRB3 knockout in aging mice via the alleviation of atrophy and fibrosis of skeletal muscles. J Cachexia Sarcopenia Muscle 11: 1104–1120

Song P, Yang S, Hua H, Zhang H, Kong Q, Wang J, Luo T, Jiang Y (2019) The regulatory protein GADD34 inhibits TRAIL-induced apoptosis via TRAF6/ERK-dependent stabilization of myeloid cell leukemia 1 in liver cancer cells. J Biol Chem 295: 5945–5955

Stoecklin G, Stubbs T, Kedersha N, Wax S, Rigby WFC, Blackwell TK, Anderson P (2004) MK2-induced tristetraprolin:14-3-3 complexes prevent stress granule association and ARE-mRNA decay. EMBO J 23: 1313–1324

Szegezdi E, Logue SE, Gorman AM, Samali A (2006) Mediators of endoplasmic reticulum stress-induced apoptosis. EMBO Rep 7: 880–885

Tamaki M, Miyashita K, Wakino S, Mitsuishi M, Hayashi K, Itoh H (2014) Chronic kidney disease reduces muscle mitochondria and exercise endurance and its exacerbation by dietary protein through inactivation of pyruvate dehydrogenase. Kidney Int 85: 1330–1339

Tanoury ZA, Piskunov A, Rochette-Egly C (2013) Vitamin A and retinoid signaling: genomic and nongenomic effects. J Lipid Res 54: 1761–1775

Thé Hd, Vivanco-Ruiz MM, Tiollais P, Stunnenberg H, Dejean A (1990) Identification of a retinoic acid responsive element in the retinoic acid receptor beta gene. Nature 343: 177–180

Vreeland AC, Yu S, Levi L, Rossetto DdB, Noy N (2014) Transcript Stabilization by the RNA-Binding Protein HuR Is Regulated by Cellular Retinoic Acid-Binding Protein 2. Mol Cell Biol 34: 2135–2146

Walter P, Ron D (2011) The unfolded protein response: from stress pathway to homeostatic regulation. Science 334: 1081–1086

Wanga Y, Pessinn JE (2013) Mechanisms for fiber-type specificity of skeletal muscle atrophy. Curr Opin Clin Nutr Metab Care 3: 243–250

Xiong G, Hindi SM, Mann AK, Gallot YS, Bohnert KR, Cavener DR, Whittemore SR, Kumar A (2017) The PERK arm of the unfolded protein response regulates satellite cell-mediated skeletal muscle regeneration. Elife 6: e22871

